# Expression Patterns of Small Interfering RNAs in Germinating Barley (*Hordeum vulgare* L.) Seeds with Age-Induced Differences in Viability

**DOI:** 10.1101/2025.07.28.667253

**Authors:** Marta Puchta-Jasińska, Paulina Bolc, Adrian Motor, Maja Boczkowska

## Abstract

Small interfering RNAs (siRNAs), a subclass of small non-coding RNAs, play crucial roles in regulating seed germination and viability through epigenetic mechanisms like RNA-directed DNA methylation (RdDM). This study presents the first comprehensive investigation of siRNA profiles linked to seed viability and germination in barley (Hordeum vulgare), utilizing a unique set of seeds from a single batch subjected to controlled long-term storage. Some seeds lost viability due to moisture exposure from unsealing, creating a natural experimental model to explore vigor effects. sRNA sequencing revealed 85,728 differentially expressed siRNAs, with distinct patterns between regenerated, high-viability, and low-viability seeds. Notably, trans-acting siRNAs (ta-siRNAs) showed peak abundance at different imbibition times depending on seed quality, suggesting dynamic regulation. Around 46% of siRNAs were 21 nucleotides, and 54% were 22 nucleotides long. Gene Ontology and degradome analyses confirmed siRNA target genes involved in vital biological processes such as cytochrome complex function, root development, cell maturation, and carbohydrate metabolism. Despite RNA degradation in low-viability seeds, siRNAs remained relatively stable, indicating their potential role in maintaining seed metabolic activity during dormancy release and germination initiation. This pioneering research uncovers novel insights into siRNA-mediated control of seed longevity and germination, highlighting the innovative use of stable, well-characterized plant material to disentangle molecular mechanisms underpinning seed vigor and germination success.

## BACKGROUND

Seed germination occurs when a dormant seed resumes metabolic activity and develops into a seedling, marking the critical transition from a seed to an independent plant [1]. It is initiated when environmental conditions such as water availability, oxygen, and optimal temperature are favorable, leading to the activation of the embryo and the emergence of the radicle through the seed coat [2]. Understanding the mechanisms and factors influencing seed germination is essential for agriculture, biodiversity conservation, and ecological restoration, as it underpins plant propagation and the survival of plant species [1]. During seed germination, metabolic and structural changes activate essential processes for successful seedling growth and development [3]. Upon water uptake (imbibition), seeds initiate repair mechanisms that reverse damage during desiccation and storage, including restoring cell membranes, organelles, DNA, and proteins [4–6]. Early germination is characterized by increased cellular respiration and the translation of stored mRNAs, followed by the onset of de novo mRNA synthesis as imbibition progresses [7–10]. As cellular hydration stabilizes, the rate of water uptake decreases, marking entry into the lag phase (phase II), during which DNA repair, mRNA transcription, and the biosynthesis of proteins involved in transcription, signaling, and metabolism occur [3, 11, 12]. Additional processes, such as cytoskeleton reconstruction, posttranslational protein modifications (e.g., ubiquitination, phosphorylation), and the synthesis of proteins necessary for root emergence, are also initiated [13–18]. The culmination of germination is marked by the emergence of the radicle from the seed coat [12, 19]. Rapid cell elongation and division drive seedling establishment in the final phase (phase III) [3, 12]. Despite significant advances, many aspects of seed germination remain incompletely understood, highlighting the need for continued research in this area. While much is known about the physiological and biochemical events that drive seed germination, the molecular regulation underlying these processes remains an active area of research. In particular, recent studies have highlighted the crucial role of gene expression regulation during germination, highlighting the involvement of small noncoding RNAs in these regulatory networks. In recent years, many researchers have focused on small noncoding RNAs. These RNAs regulate gene expression at the transcriptional and posttranslational levels[20].

The predominant classes of small RNAs (sRNAs) in plants include microRNAs (miRNAs) and small interfering RNAs (siRNAs), which differ notably in their biogenesis and functional roles [21–23]. Among these, miRNAs are the most extensively characterized, whereas the siRNA population remains poorly understood [24–27]. siRNAs are primarily generated through the cleavage of long double-stranded RNA precursors by Dicer-like (DCL) ribonucleases, which produce short duplexes that associate with ARGONAUTE (AGO) proteins to form RNA-induced silencing complexes (RISCs), which mediate gene silencing via mRNA cleavage or translational repression [28]. In plants, siRNAs play a pivotal role in epigenetic regulation through RNA-directed DNA methylation (RdDM), a mechanism crucial for seed development and genome stability. RdDM targets cytosine methylation in CG, CHG, and CHH sequence contexts, thereby modulating chromatin structure and transcriptional activity [28].

A specialized subclass of siRNAs, trans-acting siRNAs (ta-siRNAs), is produced through miRNA-directed cleavage of *TAS* gene transcripts. This process generates 21-nucleotide ta-siRNAs incorporated into AGO-RISC complexes, guiding sequence-specific cleavage or translational inhibition of target mRNAs. In *Arabidopsis thaliana*, four *TAS* gene families have been identified, which are recognized by miR173, miR390, and miR828 [29]. In association with the AGO1 or AGO7 proteins, these miRNAs regulate gene expression by inducing transcript cleavage and subsequent ta-siRNA biogenesis [30]. Like miRNAs, siRNAs mediate posttranscriptional gene silencing by guiding the degradation of complementary mRNA molecules, resulting in translational arrest [31, 32].

Overall, small RNAs regulate gene expression at both the transcriptional and posttranscriptional levels and are integral to diverse biological processes, including genome defense, plant growth and development, and responses to biotic and abiotic stresses [27, 33]. The critical role of these genes in orchestrating gene regulatory networks underscores their importance in plant adaptation and survival under fluctuating environmental conditions [27]. This regulatory complexity is exemplified in various plant species, where specific sRNAs, including miRNAs and ta-siRNAs, modulate key developmental pathways and stress responses.

For example, in cotton, fiber development is tightly regulated by two MYB family proteins whose expression correlates with that of miR828, miR858, and associated siRNAs, highlighting the interplay between transcription factors and small RNA pathways [34]. Similarly, members of the large F-box gene family, which encode substrate recognition components of SCF ubiquitin ligase complexes, are targeted by miRFBX in strawberry, where they produce siRNAs that influence fruit morphology [35]. The ARF gene family is another well-characterized target of TAS3-derived ta-siRNAs across multiple plant species, where this regulatory module governs leaf morphology, developmental transitions, root and flower architecture, embryo development, and responses to environmental stresses[36, 37]. Notably, the ARF-TAS3-ta-siRNA pathway has been demonstrated to contribute to adaptation in diverse plants, such as *Medicago truncatula*, *Lotus japonicus*, *Zea mays*, *Pyrus serotina*, and *Dimocarpus longan,* through modulation of the auxin signaling network [37]. In mosses (*Physcomitrella patens*), ta-siRNA-ARF influences sensitivity to auxin and nitrogen availability, further underscoring the evolutionary conservation and functional importance of this pathway [36]. Moreover, in *Arabidopsis*, the *TAS4* locus generates ta-siRNAs processed by miR828, which regulate genes involved in carbohydrate metabolism and stress responses, thereby linking small RNA pathways to metabolic regulation [38]. These examples illustrate the significant impact of small RNA-mediated regulation on plant development and adaptation.

Advances in high-throughput degradome sequencing have facilitated the identification of novel miRNAs and their targets, providing deeper insights into the self-regulatory networks of miRNAs and their functional roles in plant growth, development, and stress responses. Studies in *Oryza*, *Arabidopsis*, *Populus*, and *Camellia sinensis* have identified miRNAs and target genes involved in various biological processes, including terpenoid biosynthesis and stress resistance [39–42]. In maize, miRNAs associated with the regulation of seed vigor have been identified, emphasizing the importance of small RNAs in seed biology [43]. Degradome sequencing thus represents a powerful tool to validate in silico-predicted targets of small RNAs, advancing our understanding of their regulatory networks[44]. Continued research on small RNAs may expand our knowledge of their roles in seed germination and viability maintenance, ultimately contributing to crop improvement and sustainable agriculture.

Despite extensive research elucidating the roles of small interfering RNAs (siRNAs) in plant development and stress responses, their functions in dry, metabolically quiescent seeds during storage and aging remain largely unexplored. Most studies have focused on seed development, germination, or stress responses during active growth phases, leaving a significant knowledge gap regarding the molecular regulation operating in dry seeds. Although dry seeds are physiologically dormant, they undergo progressive physiological and biochemical changes during storage that can ultimately lead to loss of viability and compromised germination capacity. Emerging evidence suggests that small RNA-mediated posttranscriptional regulation may critically modulate gene expression during this latent phase, influencing pathways related to stress responses; hormone metabolism, such as abscisic acid and gibberellic acid signaling; and cellular protection mechanisms.

This study addressed the paucity of information on siRNA expression dynamics and functional roles during the postharvest dry storage of cereal seeds. We hypothesized that specific siRNAs exhibit significant expression changes during seed aging, regulating genes involved in oxidative stress responses, hormone signaling cascades, and protein turnover, which correlate with seed physiological quality and germination potential.

The primary objective was to identify and characterize siRNAs in dry barley seeds stored for various durations and investigate their potential regulatory roles in seed viability and germination. The specific goals were to (a) profile siRNA populations in dry seeds subjected to different storage periods via high-throughput sequencing technologies; (b) identify siRNAs that are differentially expressed in response to seed aging; (c) predict the target genes of these siRNAs and conduct gene ontology and pathway enrichment analyses; and (d) pinpoint candidate siRNAs that may serve as molecular biomarkers for seed longevity.

## RESULTS

### Sequencing Data Quality and Mapping

The sRNA libraries were sequenced in five separate runs. After quality filtering, low-quality reads were removed, resulting in retention rates of 35% for the Lv sample, 39% for Hv, and 45% for Rc. Approximately 3% of the raw reads corresponded to rRNA and tRNA sequences, whereas approximately 20% were aligned to the *Hordeum vulgare* reference genome (MorexV3 assembly) (Figure 1). Mapping to the reference genome identified 937,738 siRNAs (Supplementary 1). When a significance threshold of p < 0.05 was applied, 291 siRNAs were detected in Rc samples, 218 in Hv samples, and 129 in Lv samples.

**Figure 1.**
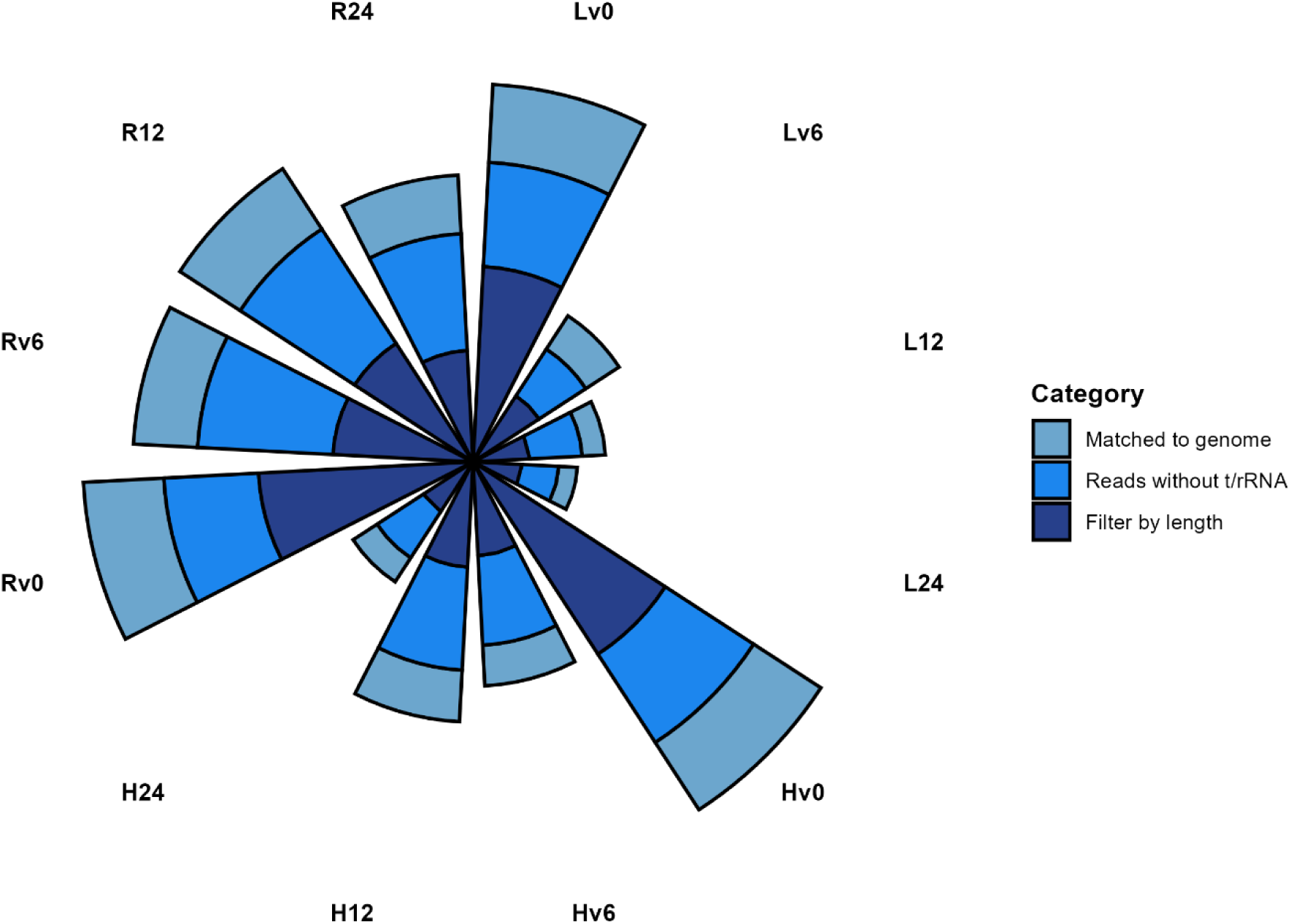
Cumulative bar graph depicting the total number of raw reads generated by small RNA sequencing (sRNA-Seq), with subsequent categorization of reads following bioinformatic processing steps, including length-based filtering, removal of tRNA and rRNA sequences, and alignment to the *Hordeum vulgare* reference genome.

### Unique siRNA Profiles Across Samples and Germination Stages

Analysis of unique siRNA sequences revealed dynamic changes during germination. In the Lv samples, 17 unique siRNAs were detected in dry seeds, increasing to 29 after 6 hours of germination, 13 after 12 hours, and 25 after 24 hours of germination. In the Hv samples, the number of unique siRNAs was 29 in dry seeds, 14 at 6 hours, 23 at 12 hours, and 15 at 24 hours. The Rc samples presented 20 unique siRNAs in dry seeds, whereas 22, 18, and 18 unique siRNAs were detected at 6, 12, and 24 h of germination, respectively (Figure 2).

**Figure 2.**
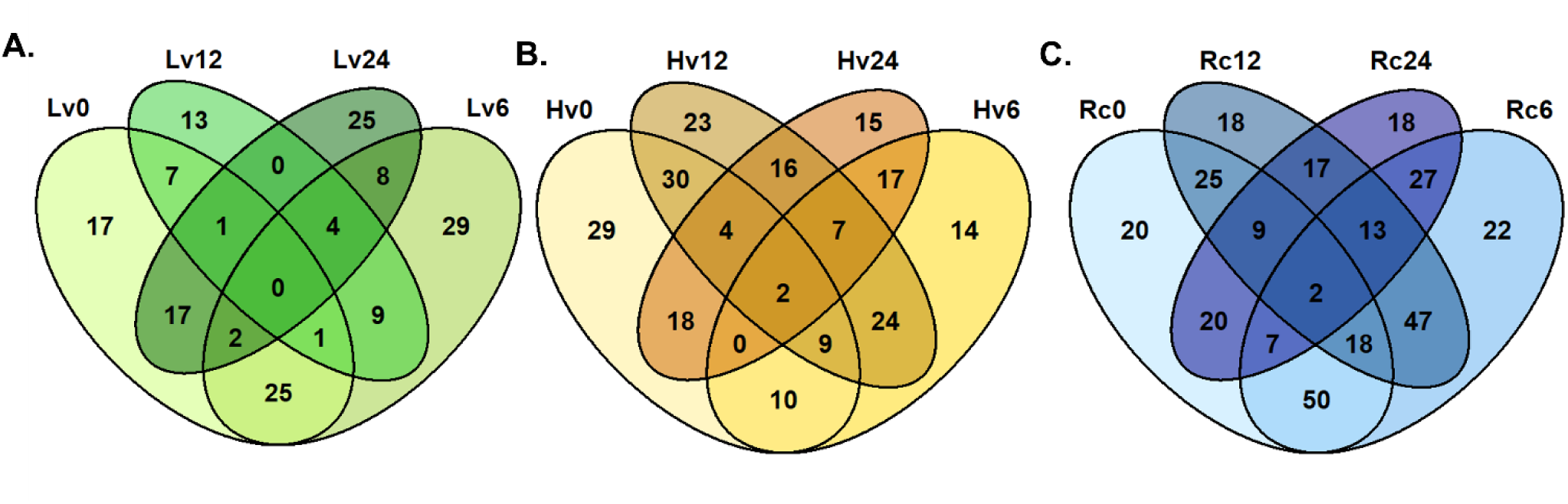
Venn diagrams illustrating the number of identified and trans-acting siRNAs (ta-siRNAs) detected during barley seed germination. (A) siRNAs and ta-siRNAs identified in low-viability (Lv) seeds at 0, 6, 12, and 24 h of germination; (B) siRNAs and ta-siRNAs identified in high-viability (Hv) seeds at corresponding germination time points; (C) siRNAs and ta-siRNAs identified in regenerated (Rc) seeds across the same germination intervals.

### siRNA Length Distribution and Expression Levels

The lengths of the identified siRNAs were predominantly 21 nt (46%) and 22 nt (54%). In the Lv dry seed sample, six siRNAs presented expression levels exceeding 2000 reads per million (RPM). At 6 h of germination (Lv6), three siRNAs (HVU_siRNA_60218, HVU_siRNA_257, and HVU_siRNA_509560) surpassed 2000 RPM. The Lv12 sample presented 25 siRNAs above this threshold, with HVU_siRNA_1501 and HVU_siRNA_2700 showing the highest expression (4444 and 4401 RPM, respectively). At 24 hours (Lv24), 23 siRNAs exceeded 2000 RPM, with the highest expression levels recorded for HVU_siRNA_114062 (5330 RPM) and HVU_siRNA_115569 (5308 RPM) (Supplementary 1).

In the Hv samples, 10 siRNAs in dry seeds exceeded 2000 RPM, with HVU_siRNA_93, HVU_siRNA_140, and HVU_siRNA_135 exhibiting the highest expression (3169, 3148, and 3137 RPM, respectively). At 6 hours, four siRNAs and six at 12 hours surpassed this threshold, whereas 12 siRNAs did so at 24 hours (Supplementary 1).

### Differential expression analysis

Differential expression analysis revealed that the greatest number of statistically significant siRNAs was detected in dry Lv seeds (Figure 3). The expression of six siRNAs (HVU_siRNA_420845, HVU_siRNA_421479, HVU_siRNA_420876, HVU_siRNA_363557,

**Figure 3.**
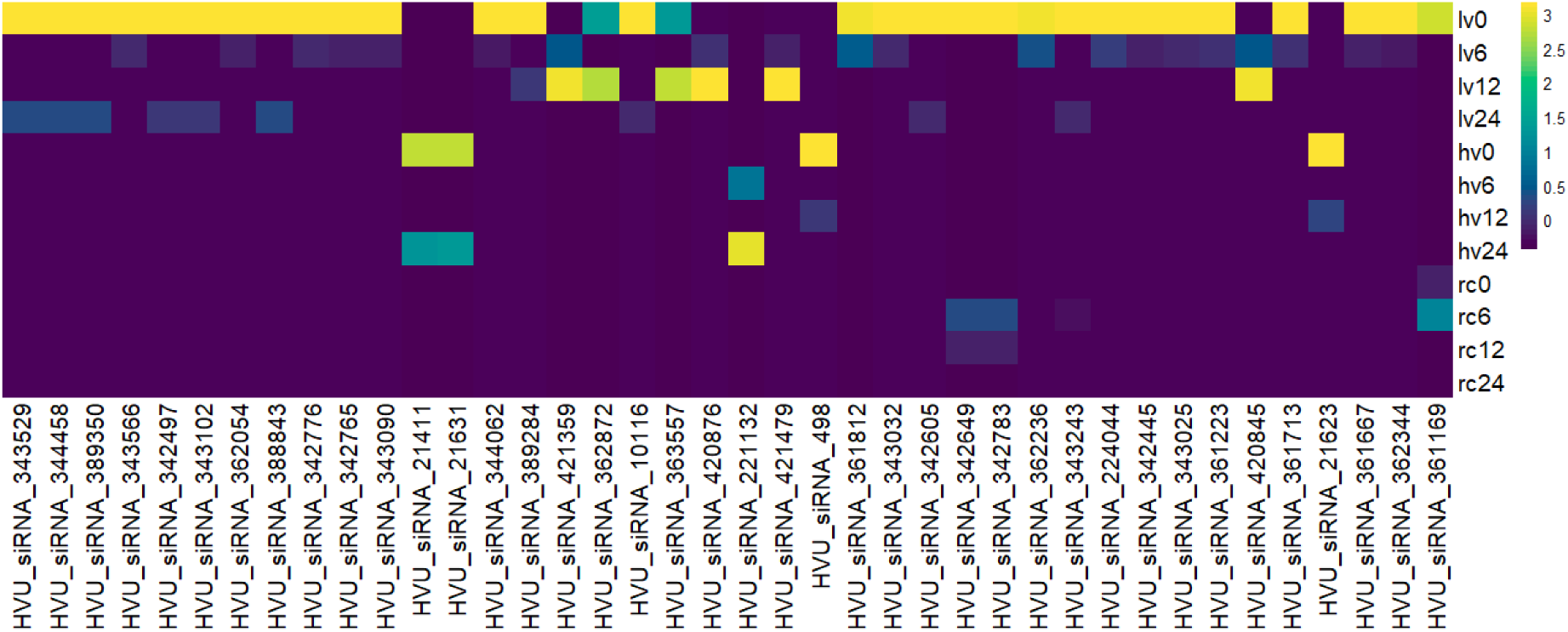
Differentially expressed siRNAs (DE-siRNAs) identified during barley seed imbibition. Changes in the expression of siRNAs with statistically significant differential expression are shown for (Hv) high-viability seeds, (Lv) low-viability seeds, and (Rc) regenerated seeds at 6, 12, and 24 hours postimbibition.

HVU_siRNA_421359, and HVU_siRNA_362872) was elevated at Lv12. HVU_siRNA_421359 and HVU_siRNA_420845 were uniquely expressed in the Lv6 and Lv12 samples. In Hv12, HVU_siRNA_498 and HVU_siRNA_21623 were highly expressed but were significantly reduced or absent in the other samples. Notably, the siRNAs with the highest differential expression were sample specific, and no differentially expressed siRNAs were detected in the Rc0 samples.

### Validation of siRNA expression by RT‒qPCR

Ten siRNAs exhibiting high expression were selected for validation via RT‒qPCR. The expression profiles obtained via RT‒qPCR were consistent with those obtained via next-generation sequencing (NGS) (Supplementary 3). For example, HVU_siRNA_186, HVU_siRNA_229922, HVU_siRNA_230263, HVU_siRNA_620, and HVU_siRNA_37149 showed strong amplification in Hv samples and were also expressed in Rc samples. HVU_siRNA_37149 was exclusive to Lv6, whereas HVU_siRNA_308723 and HVU_siRNA_308820 were amplified from the Lv24, Rc6, and Rc12 samples.

### Functional Annotation of the siRNA Targets

Gene Ontology (GO) analysis of the predicted siRNA targets revealed that in the Lv sample at 6 h of imbibition, most target genes were associated with DNA integration and metabolic processes. At 12 hours, fewer target genes presented significant GO annotations (p.adj [-log10] < 5). After 24 hours, the targets were again enriched for DNA integration and metabolism. In Hv samples, target genes were predominantly linked to chloroplast function and membrane transport across all imbibition stages. The Rc samples were enriched for DNA integration and metabolic processes at 6 hours, with sustained DNA integration-related targets at 12 and 24 hours (Supplementary 4).

### Degradome Sequencing and Target Validation

Degradome sequencing identified 481 sequences corresponding to 301 siRNAs postgermination. Among the current samples, Rc presented the highest proportion of degradome matches to siRNAs: 40.5% at 24 hours, 30.1% at 12 hours, and 10.2% at 6 hours. The Lv samples presented fewer than 5% matches at all stages, whereas the Hv samples presented 5.2% and 8.5% matches at 12 and 24 hours, respectively (Supplementary 5).

### Biological Processes Associated with siRNA Targets

Gene Ontology (GO) enrichment analysis of degradome-validated siRNA target genes revealed their involvement in various biological processes across seed viability groups and germination time points. In low-viability seeds at 6 hours postimbibition (Lv6), target genes were significantly enriched for functions such as phosphopyruvate hydratase complex activity, regulation of floral meristem growth, organ number specification, and assembly of the cytochrome b6f complex. At 12 h, the enriched processes included adenosylmethionine decarboxylase activity, nuclear membrane organization, mRNA modification and export, and polyamine metabolism. By 24 h, the siRNA targets were predominantly associated with microtubule regulation, pectin biosynthesis, cytoskeleton organization, and microtubule motor activity.

The progressive enrichment of siRNA targets involved in microtubule organization, cytoskeletal dynamics, and mRNA processing during later germination stages suggests a pivotal role for siRNAs in cellular remodeling and protein synthesis activation, both of which are critical for radicle emergence. The comparatively low representation of these processes in low-viability seeds may reflect transcriptional or posttranscriptional constraints that hinder successful germination completion.

In dry seeds, siRNAs from low-viability samples (Lv0) were linked to amino acid transport activity. High-viability (Hv) seeds at 24 hours postimbibition presented siRNA targets enriched for DNA repair complexes, the DNA replication factor C complex, DNA clamp loader activity, and salicylate 1-monooxygenase activity. Additionally, dry Hv seeds were enriched in the ketol acid reductoisomerase, adenosylmethionine decarboxylase, isoleucine metabolism, and polyamine metabolic pathways.

Regenerated (Rc) seed samples at 12 hours postimbibition were enriched for pentosyltransferase, tRNA guanosine transglycosylase, tRNA 2’-phosphotransferase, and hydrolase activities. At 6 hours, the Rc samples were enriched in purine nucleotide binding, protein folding, and ribonucleotide metabolism. The Rc dry seeds were associated with ribosome function, transcriptional regulation, catabolic process regulation, oxidoreductase activity, and mRNA decapping (Figure 4).

**Figure 4.**
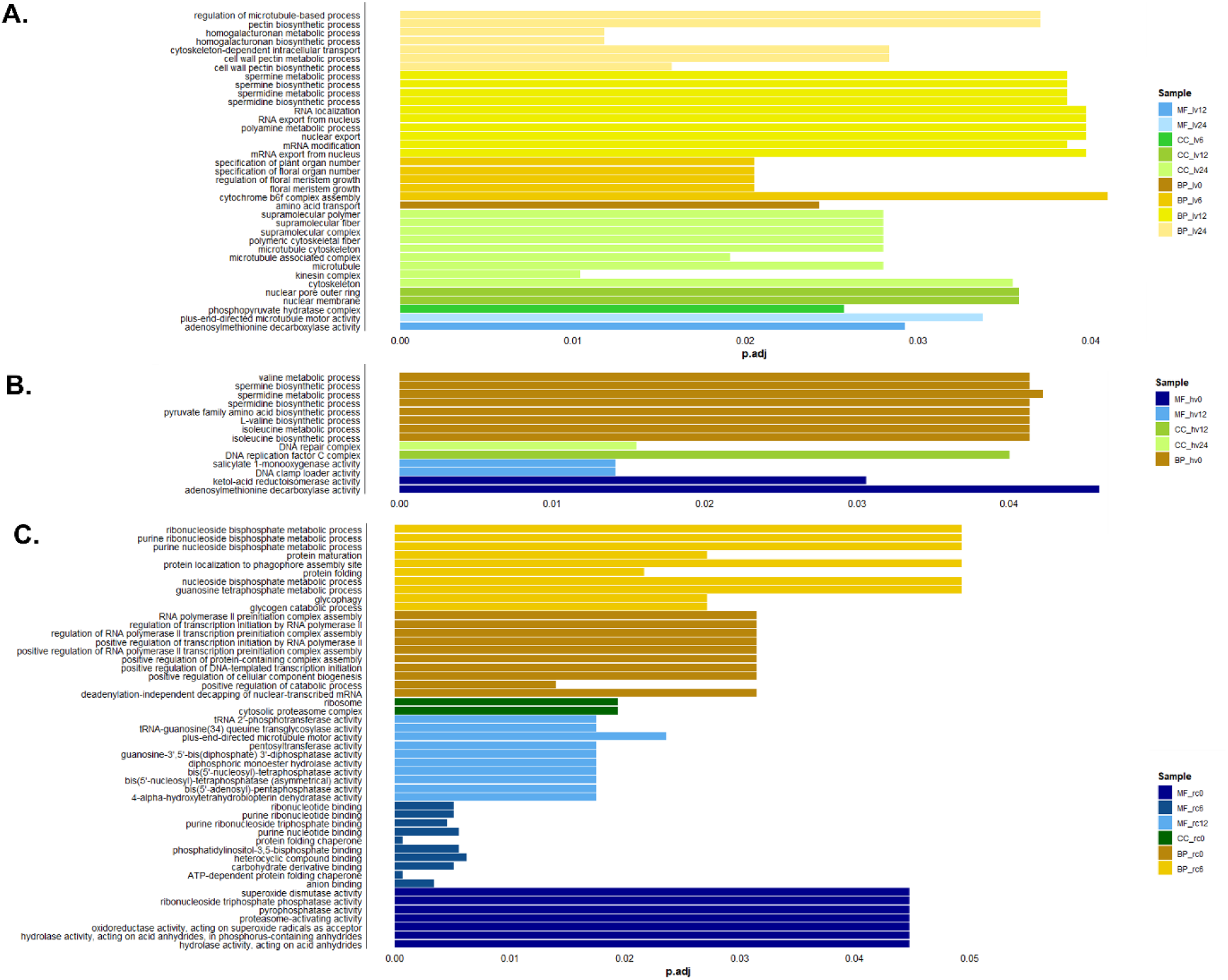
Gene Ontology (GO) enrichment analysis of degradome sequences associated with siRNAs in barley seed samples of low viability (Lv), high viability (Hv), and regenerated (Rc) groups.

### Protein function analysis of siRNA targets

Functional screening of proteins encoded by transcripts targeted by differentially expressed siRNAs revealed the involvement of DUF6598, protein disulfide isomerase, and FAR1 proteins in seed germination in Lv6 samples. Associations with the F-box, PHD, and translation initiation factors were observed in the Hv12 and Rc6 samples. At 12 hours in Lv, sample-specific DYW proteins and choline transporter-like proteins, which were also present in Rc6 and Rc24, were identified. Hexosyltransferase and kinesin proteins were linked to transcripts in the Lv24, Lv12, and Hv12 samples.

No proteins associated with degradome-validated transcripts were identified in Hv6. Hv12 transcripts related to anaphase-promoting complex proteins were detected, along with aldehyde oxygenase, transmembrane amino acid transporters, auxin efflux carriers, and MADS-box proteins, in the Rc24 and Hv24 samples. Serine/threonine kinases, WD40 repeat proteins, and tryptophan synthase were uniquely associated with Hv24 transcripts (Figure 5, Supplementary 5).

**Figure 5.**
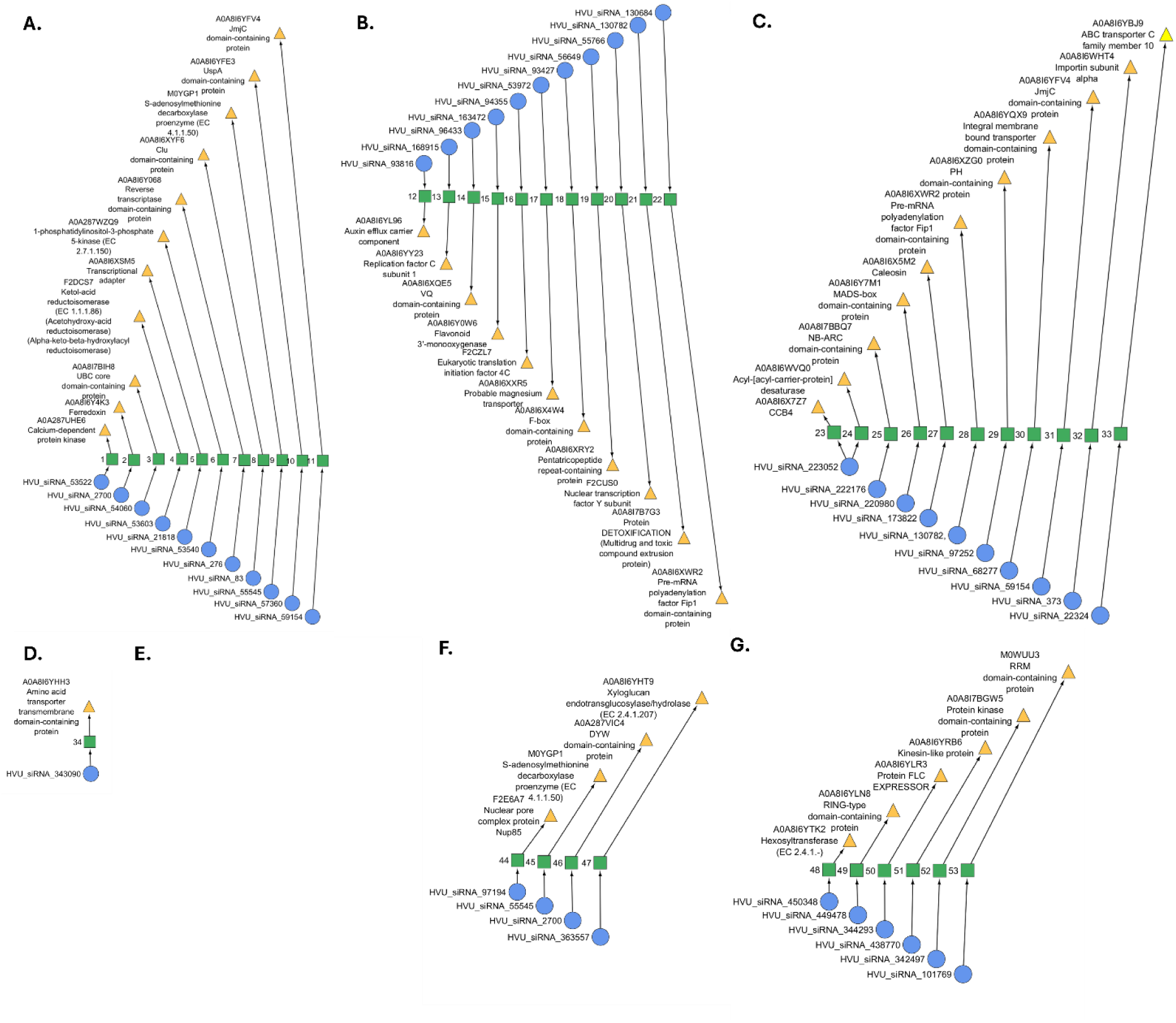
Network representation of protein annotations (yellow triangles) linked to target gene sequences (green squares) and their corresponding differentially expressed siRNAs (blue circles) across barley seed samples at various imbibition stages: (A) Hv0, (B) Hv12, (C) Hv24, (D) Lv0, (E) Lv6, (F) Lv12, and (G) Lv24. Protein identifiers correspond to entries in the UniProt database. Detailed information on the target gene sequences is provided in Supplementary 5.

## DISCUSSION

### Role of siRNAs in germination

Small noncoding RNAs (sRNAs) are critical in plant development and response to environmental stresses [45, 46]. While extensive research has elucidated the functions of microRNAs (miRNAs) in seed development and germination, the contributions of small interfering RNAs (siRNAs) to these processes remain underexplored [47, 48]. Notably, siRNA involvement has been documented in the male germline, where RNA-directed DNA methylation (RdDM) activity is markedly reduced concomitant with increased siRNA expression [49]. In olive (*Olea europaea*), the accumulation of 24-nucleotide sRNA sequences during the extended juvenile phase suggests a potential role for siRNAs in maintaining seed dormancy and regulating aging [50]. The germination process is governed by a complex regulatory network in which sRNAs play a key role as modulators [51, 52].

Endogenous siRNAs mediate gene silencing through mechanisms including posttranscriptional gene silencing and chromatin remodeling, thereby inactivating homologous sequences at multiple regulatory levels [53, 54]. Our results revealed distinct sRNA expression profiles in seed samples whose viability varied during the initial 24 h of germination. These differential expression patterns underscore the dynamic regulatory capacity of siRNAs in modulating seed viability and germination timing. In this study, we identified 1,324 small interfering RNAs (siRNAs) and 217 trans-acting siRNAs (ta-siRNAs) with temporally dynamic expression profiles during the first 24 hours of imbibition in barley (*Hordeum vulgare*) seeds with distinct viability levels. Comparative analysis revealed statistically significant differential expression patterns between low-viability (Lv) and high-viability (Hv) seed cohorts, suggesting that siRNA-mediated regulatory mechanisms are integral to the transcriptional reprogramming required to transition from dormancy to active germination. These findings underscore the potential role of siRNAs in modulating stress adaptation, epigenetic regulation, and metabolic activation during early germination stages.

This study utilized unique plant material from a single seed lot subjected to long-term storage, effectively minimizing the variability attributable to environmental factors. During storage, one subset of seeds was exposed to increased moisture following unsealing the container, resulting in a pronounced decline in viability. This natural aging process provides an opportunity to investigate the influence of seed vigor on the germination dynamics of seeds with inherently different viability levels.

### Stability of siRNA Fractions in Seeds with Varying Viability During Germination

The size distribution of small RNAs (sRNAs) is highly conserved across angiosperms[50]. However, distinct differences exist among plant clades; for example, gymnosperm seeds predominantly contain 24-nucleotide (nt) siRNAs, whereas in other gymnosperms, 24-nt siRNAs are nearly absent, highlighting clade-specific siRNA profiles[55]. In barley cultivars Golden Promise and Pallas, 23–24 nt sRNAs have been implicated in repressive chromatin modifications and genome stability, whereas 20–21 nt sRNAs contribute to varietal specificity[56]. Chen et al. (2009) demonstrated that 21- and 22-nt siRNAs associate with ARGONAUTE1 (AGO1) to mediate mRNA cleavage, whereas 23- and 24-nt siRNAs interact with AGO4 and AGO6 to promote DNA methylation at asymmetric CHH and H3K9 sites, thereby maintaining genomic integrity[24]. In the present study, 46% of the identified siRNAs were 21 nt in length, and 54% were 22 nt in length, suggesting that their primary function may involve target mRNA cleavage, which is consistent with previous reports.

Seed viability loss is accompanied by macromolecular damage affecting DNA, RNA, and proteins [57]. RNA molecules are particularly susceptible to degradation during seed aging, with total RNA quantity and integrity decreasing as viability decreases [58–60]. This degradation is heterogeneous across RNA fractions; notably, small RNAs, including miRNAs and siRNAs, are relatively stable and do not degrade in proportion to the loss of seed viability and RNA integrity [61]. siRNAs function in concert with miRNAs within the RNA-induced silencing complex (RISC), whose protein components exhibit increased expression during germination [62]. Thus, it is plausible that siRNA levels remain relatively stable in stored seeds, with variations emerging primarily during subsequent imbibition stages.

Analysis of siRNA abundance across germination stages revealed that variability was correlated with seed viability. Compared with low-viability seeds, regenerated seeds presented more identified siRNAs, suggesting that siRNAs play a role in metabolic processes critical for dormancy release and germination initiation. In low-viability seeds, siRNA numbers increased progressively during imbibition, potentially reflecting delayed germination onset and greater degradation of stored mRNAs. Similar trends have been observed for miRNAs, where samples with reduced viability presented increased miRNA abundance at later germination stages [23].

In rice, Meijer et al. (2022) reported that sRNA loci are three to seven times more abundant in embryos than in other tissues, although their expression decreases with increasing grain age [55]. Rapid cell division in developing embryos, such as those of Arabidopsis, has been linked to increased expression of 24-nt transposable element (TE)-derived small interfering RNAs (siRNAs) [56]. Studies in *Brassica rapa* identified 24-nt siRNAs transcribed from protein-coding regions, whereas analyses in Chinese cabbage demonstrated that 20–24 nt siRNA abundance varies with the duration of heat stress[63]. Additionally, in *Caenorhabditis elegans*, 21-nt siRNAs exhibit high complementarity to mRNAs [64]. These findings and our data suggest that the observed differences in siRNA content across germination stages reflect progressive aging processes and variations in germination capacity.

Germination is regulated by a complex network of molecular mechanisms, with sRNAs playing integral roles [49]. Our study revealed distinct siRNA expression patterns in seeds with varying viability during the first 24 h of imbibition, highlighting the dynamic regulation of protein expression by siRNA-mediated mechanisms. Unlike miRNAs, conserved siRNAs standard for all samples were not detected, implying that siRNAs may exert specialized control over proteins essential for successful germination.

### Functional Analysis of siRNA-Regulated Genes in Seeds with Varying Viability

The process of germination of dry seeds requires the resumption of metabolism and initiates exchanges at the transcription level.

The germination of dry seeds necessitates the resumption of metabolic activity and initiates extensive transcriptional reprogramming. Target genes identified through in silico prediction were validated via degradome sequencing to elucidate the functions of siRNA-regulated genes. This analysis yielded 481 degradome sequences corresponding to 301 siRNAs, representing approximately 10% of all in silico-predicted targets. Notably, over 80% of these sequences were detected in regenerated seeds, whereas only 13% and 5.6% were detected in long-term stored seeds with high and low viability, respectively. These findings suggest a positive correlation between the abundance of siRNA-targeted transcripts and the retention of germination capacity. In a study of Arabidopsis mutants, 1,247 degradome sequences were identified. Comparative studies in *Arabidopsis* have reported similar approaches, with degradome sequencing confirming targets for numerous siRNAs, underscoring the utility of this method in functional validation [39, 65].

#### Functional Categories of siRNA-Regulated Proteins in High-Viability Seeds

Functional analysis of proteins associated with high seed viability revealed their involvement in membrane trafficking processes linked to the AP5 adaptor protein complex and ribonuclease activity. Notably, proteins from the MADS-box family, histone deacetylases, peroxidases, tryptophan synthases, integral membrane transporters, detoxification-related proteins, NB-ARC domain-containing proteins, F-box proteins, and domains of unknown function (DUF) were prominently represented.

The data obtained via degradome analysis demonstrated that siRNAs in highly viable seed samples (Hv and Rc) targeted a variety of regulatory proteins associated with development, oxidative stress, translational reprogramming and transcriptional activation (Supplementary 6). In highly viable seeds, the presence of NB-ARC domain-containing proteins, which function as immune receptors and sensors of molecular damage, is particularly significant. Repressing these proteins via RNA interference may be crucial for inhibiting excessive stress responses in regenerating cells. NB-ARC domain-containing proteins play essential roles in plant development, including processes relevant to seed germination and early growth stages. However, specific, direct studies on seed germination are limited. The NB-ARC domain (nucleotide-binding adaptor shared by APAF-1, R proteins, and CED-4) is a conserved nucleotide-binding domain that functions as a molecular switch by binding and hydrolyzing ATP or ADP, thereby regulating protein activity and downstream signaling pathways that are vital for plant development and defense [66]. Furthermore, studies have identified siRNAs that target domains associated with PHD-finger, F-box, MADS-box, and S/T kinases in highly viable seeds, which play pivotal roles in cell cycle control, protein ubiquitination, and transcriptional regulation. These proteins play crucial roles in regulating phase transitions within the cell cycle and activating transcription processes that govern the expression of developmental genes [2, 67–69]. The repression or modulation of siRNAs may serve as a mechanism for the temporal control of metabolic reprogramming in germination-recovery seeds.

Although DUF proteins remain poorly characterized, they are hypothesized to participate in stress response and repair pathways critical for maintaining seed viability. These proteins belong to the serine esterase superfamily, which has been implicated in defense signaling against herbivorous insects, as demonstrated by Ortiza et al. [87]. Gao et al. further classified DUF proteins in *Oryza sativa*, *Zea mays*, and *Hordeum vulgare* as key components of pathogen response mechanisms [70]. Oligosaccharides play essential roles in maintaining cell wall integrity and function, with numerous DUF family genes involved in polysaccharide biosynthesis and modification via glycosyltransferase activity. Specifically, DUF266 domain-containing proteins contribute to the biosynthesis of cellulose, a major structural component of the plant cell wall. The functional disruption of DUF genes in rice has been shown to reduce cellulose content, underscoring their importance in cell wall formation [71]. Our results further corroborate these findings by demonstrating that in barley seeds with high viability, DUF proteins are actively expressed and likely contribute to preserving cell wall integrity and orchestrating stress response pathways, thereby supporting seed longevity and successful germination.

Several siRNAs have also targeted transcripts linked to abscisic acid-mediated germination inhibition pathways. Abscisic acid (ABA) inhibits seed germination primarily by preventing embryo cell wall loosening and restricting water uptake, thereby limiting radicle emergence and cell expansion during germination[72, 73]. This inhibition is mediated through the suppression of genes encoding cell wall biosynthetic and remodeling enzymes, including expansins, pectin esterases, xyloglucan endotransglycosylases, cellulose synthases, and extensins, which are essential for cell wall loosening and embryo growth [74]. The reversible nature of the effect of ABA allows germination to proceed upon its removal, highlighting its regulatory role in maintaining seed dormancy under unfavorable conditions [73, 75].

The overexpression of DUF4228 in *Medicago sativa* enhances ABA-responsive gene expression, leading to increased ABA accumulation and reduced seed germination rates, which suggests a link between specific protein families and ABA-mediated germination control[76]. F-box domain proteins, which function in proteasomal ubiquitination, also participate in ABA-regulated germination. In rice, OsFbox352 expression is upregulated by ABA and downregulated by glucose, suggesting that F-box proteins mediate glucose-dependent modulation of ABA metabolism to inhibit germination [86]. Similarly, in *Arabidopsis*, F-box proteins such as CTG10 (cold temperature germinating 10) promote germination by destabilizing PIF1, a negative regulator of germination, under light conditions. The overexpression of CTG10 reduces sensitivity to gibberellin biosynthesis inhibitors, whereas mutants exhibit hypersensitivity [77]. Members of the F-box auxin signaling family (AFB1 and AFB5) negatively regulate germination via the auxin pathway in maternal tissues, with mutants exhibiting accelerated germination compared with the wild type [78]. These findings underscore the multifaceted role of ABA in inhibiting germination through the modulation of cell wall dynamics, hormonal signaling pathways, and protein degradation systems, thereby integrating environmental and metabolic cues to regulate seed dormancy and germination timing.

NB-ARC domain-containing proteins play critical roles in pathogen recognition, activation of defense signaling cascades, and regulation of cellular development. In highly viable seeds, repression of these targets may reduce unnecessary activation of defense responses, facilitating resource allocation toward growth. This balance may be critical for successful germination and early seedling development. These proteins catalyze ADP hydrolysis through their ARC domain, thereby modulating plant developmental processes [79]. Recent studies have revealed that NB-ARC proteins are also integral components of innate immunity and programmed cell death pathways in metazoans[80]. Notably, NB-ARC genes, such as human APAF-1 and *Caenorhabditis elegans* CED-4, share homology with key regulators of apoptosis, underscoring their conserved role in regulating cell death across kingdoms [81]. Within plants, members of the NB-ARC gene family, such as *Arabidopsis* RPP1A, contribute to disease resistance against virulent strains of *Hyaloperonospora parasitica* but may also reduce plant growth [82]. The expression of the TIR-NB-ARC-LRR gene VpTNL1 in *Arabidopsis* has been associated with phenotypic variation ranging from wild-type to dwarf morphologies [83]. Similarly, the maize NB-ARC protein AhRAF4 is correlated with developmental morphological changes [84]. A prominent member of this family, the mother of FT and TFL1 (MFT), regulates seed germination by modulating hormone signaling pathways involving abscisic acid (ABA) and gibberellins (GA) [72]. MFT is a key regulator of seed sensitivity to ABA, influencing the balance between germination initiation and dormancy maintenance [85, 86]. During seed germination, intensive amino acid biosynthesis occurs, concomitant with the hydrolysis of storage proteins. This process leads to a marked increase in free amino acids, including essential amino acids such as tryptophan, as observed in soybean seed embryos [87].

Our results also revealed that siRNAs are involved in the metabolism of tryptophan. Exogenous application of tryptophan to seeds enhances amylase activity during early seedling development, facilitating increased sugar availability during subsequent metabolic stages and promoting the degradation of reserve starch. Amylases and glucosidases hydrolyze carbohydrate reserves in seeds into glucose, which serves as an energy source or is transported to the embryo axis as sucrose [88–90]. Hussain et al. reported that tryptophan treatment of sunflower seeds significantly improved germination rates, root and shoot lengths, and biomass accumulation [91].

Finally, siRNAs target histone deacetylases (HDACs), pivotal regulators of seed development, dormancy, and aging through epigenetic mechanisms [92]. Inhibition or mutation of HDAC activity results in histone hyperacetylation, leading to altered gene expression patterns that negatively impact seed viability and longevity [93]. In *Arabidopsis thaliana*, HDA19 interacts with HSL1 to repress seed maturation genes during germination. Loss of HDA19 function leads to aberrant gene expression, characterized by increased acetylation of histones H3 and H4 (H3ac, H4ac) and the activation of histone modifications such as H3K4me3[94]. Additionally, the plant-specific HDACs HD2A and HD2B suppress the expression of the delay of germination 1 (DOG1) gene, a key regulator of seed dormancy. Loss of HD2A and HD2B increases histone acetylation at DOG1 loci, increasing dormancy and potentially extending seed viability by delaying germination under stress conditions[95].

Germination requires the transition of dry seeds from quiescence to active metabolism, accompanied by extensive transcriptional reprogramming. The functional analysis of proteins correlated with high seed viability revealed their involvement in membrane transport, transcriptional regulation, and cell differentiation processes. These findings suggest that initiating metabolic activity during germination involves cell wall permeability and hormonal regulation modifications, which are essential for breaking seed dormancy.

#### Functional Categories of siRNA-Regulated Proteins in Low-Viability Seeds

Functional analysis of siRNAs in seeds with low viability (Lv) revealed several differentially expressed siRNAs that target transcripts encoding proteins involved in stress responses, mRNA processing, and metabolic regulation (Supplementary 5). FAR1 family proteins, PHD finger domain proteins, protein disulfide isomerases, choline transporters, DYW domain-containing pentatricopeptide repeat (PPR) proteins, and translation initiation factor 4C. These small interfering RNA (siRNA) targets play functional roles in mRNA modification, membrane-associated activities, sugar biosynthesis, and isomerase activity. The correlations observed in low-viability seeds suggest the initiation but incomplete execution of metabolic processes. Previous studies on dry-stored barley seeds have similarly demonstrated the onset of metabolic activities during storage, with transcript damage in low-viability seeds likely impeding proper germination completion [96].

Our data indicate that in the Lv6 and Lv12 samples, siRNAs were enriched for targets associated with organelle-localized RNA editing and chromatin regulation. FAR1 family proteins function as far-red light-responsive transcription factors that modulate seed dormancy and germination by regulating sensitivity to abscisic acid (ABA), a hormone that inhibits germination. Maize mutants deficient in FHY3 and FAR1 exhibit reduced ABA sensitivity during germination, indicating a role in promoting germination under favorable conditions [97]. Additionally, FAR1 proteins contribute to accelerated leaf senescence by influencing chlorophyll synthesis and photosystem activity in *Arabidopsis*[98].

DYW domain-containing proteins, which are involved in RNA editing in chloroplasts and mitochondria, were also targeted, indicating potential disruption of organelle gene expression, which is essential for energy metabolism during germination. DYW domain proteins, pentatricopeptide repeat (PPR) family members, mediate RNA editing within plant organelles [99]. Although not directly linked to seed aging, their role in RNA editing may affect the expression of genes critical for seed vigor and stress responses. Proper RNA editing in mitochondria and chloroplasts is crucial for maintaining energy metabolism, a key factor in determining seed germination and plant growth. Disruption of these processes in aging seeds suggests that DYW proteins may be vital for sustaining seed viability [100].

PHD finger proteins are among the siRNA targets in Lv samples; they modulate the expression of genes related to gibberellin (GA) and ABA biosynthesis and catabolism during germination [101]. For example, PIL5 (PIF1), a phytochrome-associated protein, activates the SOM and DAG1 genes, inhibiting GA biosynthesis and increasing ABA biosynthesis, thereby suppressing germination [102, 103].

Translation initiation factors, which are targeted by siRNAs in low-viability seeds, are crucial during the early germination stages, enabling the selective translation of stored mRNAs necessary for energy metabolism, detoxification, and stress responses [10, 104]. Reduced translational capacity in aged seeds, often due to oxidative damage to stored mRNAs, is correlated with diminished germination potential. Additionally, choline transporter-like proteins, associated with phospholipid biosynthesis and membrane integrity, were identified as siRNA targets in Lv samples. Choline also participates in signal transduction during stress responses. Its protective role against oxidative damage may indirectly support protein translation by preserving cellular structure and function [105, 106].

These results suggest that although metabolic processes are reactivated in low-viability seeds, damage to transcripts and compromised molecular functions impede the successful completion of germination. This observation is consistent with previous studies on barley seeds, which demonstrated that while metabolic activation occurs during storage, the loss of transcript integrity has an adverse effect on germination outcomes [96].

From an application perspective, the differential abundance and stability of specific siRNAs across seed viability gradients present promising opportunities for developing molecular biomarkers to assess seed quality. The implementation of siRNA-based biomarker assays could complement conventional germination tests by providing more rapid and potentially more accurate predictions of seed viability, thereby benefiting seed conservation efforts and agricultural practices. Nonetheless, the practical utility of such molecular tools necessitates rigorous validation of candidate siRNAs across diverse genotypes and storage environments to ensure robustness and generalizability.

#### Key Processes Regulated by siRNAs During Seed Germination across Viability Levels

Our results indicate that siRNAs play crucial roles in initiating metabolic resumption and exerting regulatory control during seed germination. Functional analyses revealed the involvement of histone deacetylases (HDACs) and mRNA metabolism-related processes predominantly in seeds with high viability [107]. Histone acetylation and deacetylation are critical epigenetic modifications that regulate gene expression in eukaryotes. HDACs catalyze the removal of acetyl groups from lysine residues on histone tails, leading to chromatin condensation and transcriptional repression. This mechanism plays a pivotal role in breaking seed dormancy by tightly regulating chromatin states. Loss or perturbation of HDAC function induces epigenetic instability and developmental abnormalities, ultimately leading to seed senescence. Studies in yeast have demonstrated that inactivation of HDAC complexes extends the cellular lifespan and enhances stress resistance by modulating trehalose metabolism, suggesting a conserved role for HDACs in aging regulation across species [108]. In humans, HDAC2-containing complexes have been implicated in aging processes, underscoring the broad biological significance of HDACs in longevity [107, 109].

In *Arabidopsis thaliana*, loss-of-function mutations in HDA6 reduce seed germination rates under abscisic acid (ABA) and salt stress conditions [110]. Furthermore, HDA19 interacts with the SIN3 complex, which associates with ERF7 to regulate ABA-responsive gene expression during ABA treatment and drought stress [111]. Recent findings indicate that histone deacetylation complex 1 (HDC1) physically interacts with both HDA6 and HDA19, increasing seed germination under stress and ABA exposure in *Arabidopsis* [110]. In rice (*Oryza sativa*), the overexpression of HDT701 increases susceptibility to pathogens while conferring increased tolerance to salt and osmotic stress [94, 112]. Imbalances in histone acetylation dynamics disrupt mitochondrial function and reduce ATP synthesis, leading to irreversible cellular damage. Notably, aged rice seeds exhibit decreased expression of GA20ox genes, which are essential for gibberellin biosynthesis; this downregulation correlates with the accumulation of oxidative damage and a loss of seed viability [113]. Han et al. (2023) demonstrated that HD2A- and HD2B-mediated repression of the delay of germination 1 (DOG1) gene is crucial during seed maturation and storage in Arabidopsis. DOG1 functions as a conserved regulator of dormancy by modulating gibberellin metabolism [95].

Pharmacological inhibition of HDAC activity by trichostatin A (TSA) delays germination in maize [114]. Conversely, *Arabidopsis* HDA9 mutants display reduced dormancy, increased germination rates, and increased resistance to artificial aging [115]. These findings, which are supported by previous literature, collectively underscore the essential role of HDACs in maintaining seed viability and facilitating proper germination. By modulating chromatin condensation, HDACs likely influence the transcriptional activity of seed-stored mRNAs in dry seeds, thereby promoting or repressing germination processes.

### Model of Barley Seed Viability Regulation by siRNA: From Precise Control to Progressive Deregulation

This study proposes a comprehensive model that elucidates the regulatory role of small interfering RNAs (siRNAs) in barley seed viability on the basis of differential expression profiles and target gene analyses in seeds with varying germination capacities (Figure 6). Our model delineates a fundamental dichotomy in siRNA function corresponding to seed viability status. In high-viability (Hv) seeds, siRNAs act as precise modulators that maintain cellular homeostasis by fine-tuning stress response pathways; regulating epigenetic and developmental gene networks, including histone deacetylases and MADS-box transcription factors; and safeguarding genomic integrity through targeting DNA repair and replication machinery components. Such a well-coordinated system can facilitate a successful transition from dormant to active metabolism, ensuring germination and growth. However, in low-viability (LV) seeds, siRNA profiles exhibit progressive deregulation, characterized by increased siRNA diversity resulting from uncontrolled mRNA degradation. Here, siRNAs aberrantly target essential transcripts involved in mitochondrial and chloroplast RNA editing (notably DYW domain-containing proteins), translation initiation factors, and structural components such as cytoskeletal and membrane-associated proteins.

**Figure 6.**
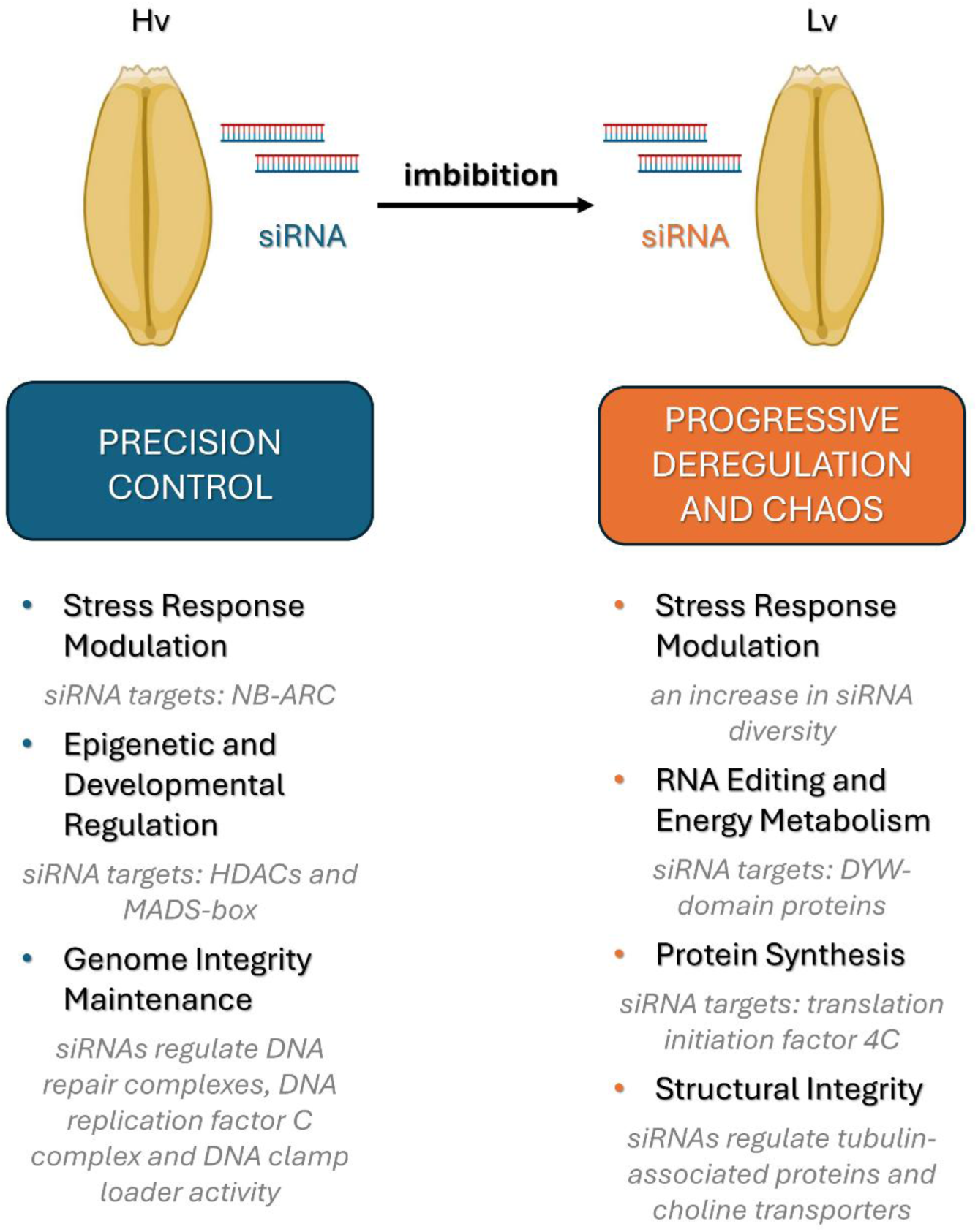
The proposed model of siRNA regulation of barley seed viability is derived from an integrative analysis of siRNA expression profiles and degradome data in barley seeds exhibiting low viability (Lv) and high viability (Hv).

This mistargeting disrupts energy metabolism, protein synthesis, and cellular architecture, thereby exacerbating molecular damage and contributing to systemic cellular failure. These findings reveal a dualistic role for siRNAs in determining seed viability. siRNAs transition from guardians of molecular homeostasis to agents of cellular disintegration. These findings provide new insights into the molecular mechanisms underlying seed aging and loss of viability.

### Study limitations

We are aware that this study has several limitations. The functional roles of the identified siRNAs were predominantly inferred through in silico target prediction and degradome sequencing, which lack direct genetic validation or functional characterization via transgenic approaches. Consequently, it remains uncertain whether these siRNAs actively mediate the regulation of germination or merely represent secondary molecular changes associated with seed aging. Additionally, the limited matching between predicted siRNA targets and degradome-approved transcripts suggests that more accurate target prediction tools and/or increased sequencing depth are necessary to ensure that siRNA targets are identified correctly and comprehensively.

### Future Directions

These findings advance the current understanding of siRNA dynamics in barley seeds exhibiting differential viability and highlight several promising avenues for future research. While the roles of microRNAs in seed dormancy, stress responses, and longevity have been relatively well characterized, a more detailed investigation into the distinct and potentially overlapping functions of small interfering RNAs is warranted. Functional validation of key differentially expressed siRNAs identified in this study should be pursued through transgenic approaches, including overexpression, RNA interference, or CRISPR/Cas-mediated modulation of siRNA-generating loci. Furthermore, the generation and phenotypic characterization of barley lines deficient in core siRNA pathway components—such as Dicer-like proteins, RNA-dependent RNA polymerases, or specific ARGONAUTE isoforms—will provide critical insights into whether altered siRNA biogenesis influences seed aging, germination capacity, or transcriptome integrity under prolonged storage conditions. Integrative analyses simultaneously profiling miRNAs, siRNAs, and long noncoding RNAs within the same seed samples could elucidate whether these regulatory pathways operate synergistically, antagonistically, or independently during the transition from dormancy to active metabolism. Given that many siRNAs mediate gene silencing via RNA-directed DNA methylation (RdDM), whole-genome methylome profiling could be used to determine whether siRNA expression patterns correlate with epigenomic reprogramming events associated with germination or seed aging. Comparative small interfering RNA (siRNA) profiling across multiple cereal species and diverse storage environments would facilitate the identification of evolutionarily conserved and species-specific regulatory motifs, thereby enabling the development of universal molecular biomarkers for seed viability. Moreover, integrating siRNA expression data with transcriptomic, proteomic, and metabolomic datasets could reveal upstream regulators and downstream physiological processes governed by siRNA-mediated control. Such comprehensive, multiomics approaches can potentially establish small interfering RNAs (siRNAs) as predictive indicators of seed viability. This study provides a valuable molecular complement to conventional germination tests in gene bank management and crop improvement programs.

## CONCLUSIONS

Our integrative analysis of siRNA expression and target profiling in barley seeds of differing viability levels reveals critical regulatory roles for small RNAs in modulating germination potential. In highly viable seeds, siRNAs selectively target transcripts associated with transcriptional regulation, membrane trafficking, and cell cycle control, suggesting a role in fine-tuning developmental transitions during early germination. Conversely, in low-viability seeds, siRNAs were enriched for targets involved in stress responses, RNA processing, and translation initiation, indicating disrupted but initiated metabolic reprogramming. The repression of NB-ARC domain proteins, PHD finger transcription factors, and DUF-containing proteins in viable seeds implies strategic silencing of stress or senescence pathways to ensure successful germination. In contrast, the downregulation of DYW domain proteins and translation factors in aged seeds likely impedes organelle function and protein synthesis. These diverse siRNA profiles emphasize the interaction between regulatory RNA pathways and the physiological state of seeds. Our findings support the hypothesis that siRNAs serve as both posttranscriptional regulators and potential biomarkers of seed viability. siRNA-based diagnostic methods could increase the accuracy of seed viability tests in gene banks. Future efforts should focus on validating these molecular markers across various genotypes and storage conditions to enable their broader application.

## METHODS

### Plant material

Seeds of the Damazy cultivar of barley, harvested in 1972, were used in this study. After being harvested and dried, the seeds were stored in air-filled, hermetically sealed containers at room temperature. The seeds had more than 95% viability and a moisture content of 2.96%. Upon reassessment in 2015, the viability of the seeds decreased to 2%, while the moisture content increased to 12.5%. For reference, a separate batch of barley seeds was kept under the same conditions for the same duration. These seeds showed 99.3% viability and a moisture content of 6.32% in 2015. They were regenerated in a field trial in 2019 and used as control samples. Puchta et al. previously described a comprehensive description of these plant materials[61].

For the experimental analysis, seeds from each sample were germinated in 60-mm-diameter Petri dishes lined with moist cotton. Embryos and scutellum were collected from dry seeds and at three different time points during germination (after 6, 12, and 24 hours). Each time point was examined in three independent biological replicates, each consisting of 25 seeds.

### sRNA extraction and construction of sRNA libraries

Small RNA (sRNA) was extracted from 36 samples via a commercial microRNA isolation kit from A&A Biotechnology (catalog number 035--100). For each sample of 25 embryos, the tissue was ground to a fine powder in a mortar under liquid nitrogen to preserve RNA integrity. The isolated sRNAs were then used to construct sRNA libraries following the protocol described by Puchta et al. (2021)[61]. Size selection was performed via automated Pippin Prep electrophoresis to ensure enrichment of sRNA fragments of the desired length. Thirty-six sRNA libraries were generated and sequenced on an Illumina MiSeq platform via Reagent Kit v3 (150 cycles) to obtain 51 bp single-end reads.

### Degradome sequencing

Total RNA was extracted via TRIzol reagent (Invitrogen) according to the manufacturer’s protocol. Polyadenylated [poly(A)] RNA was subsequently isolated from the total RNA fraction via magnetic bead-based purification. To capture the 3′ cleavage products of mRNA, which possess a 5′ monophosphate, RNA ligase was employed to ligate 5′ adapters specifically to the 5′ ends of these fragments. Following adapter ligation, the fragments were purified via magnetic beads. Complementary DNA (cDNA) synthesis was performed using 3′ and 5′ cDNA primers, followed by a second purification step with magnetic beads. The PCR products were subjected to enzymatic digestion with the MmeI endonuclease, and duplex adaptors were ligated to the resulting fragments. The ligation products were size-selected and purified via electrophoresis on a high-resolution 4% MetaPhor agarose gel. PCR amplification was subsequently carried out via primers compatible with the Illumina sequencing platform. Finally, paired-end sequencing (150 bp reads) was performed on an Illumina NovaSeq system. The detailed degradome library construction protocol is described in Puchta-Jasińska et al.[116].

### Bioinformatic analysis

The quality of the raw sequencing reads was evaluated via FastQC software. Low-quality reads (Phred score < 30) and adapter sequences were removed via the UEA Small RNA Workbench [117]. The high-quality reads were aligned to the *Hordeum vulgare* reference genome (MorexV3_pseudomolecules_assembly), retrieved from the Ensembl Plants database (Release 57, accessed 17 February 2023). SiRNA and trans-acting siRNA (ta-siRNA) loci were identified via the SiLoCo and tasiRNA tools integrated within the UEA Small RNA Workbench [118]. Custom Python scripts were developed to facilitate the identification of siRNA sequences [39]. Novel gene prediction was conducted in silico via the PsRobot tool [119].

The degradome sequencing data were subjected to quality control and mapped to the same reference genome assembly. Target validation was performed in silico via the CleaveLand 4.5 pipeline [120].

Quantitative differential expression analyses were conducted via DESeq2, implemented in the SARTools R package, where statistical significance was assessed via Fisher’s exact test to identify genes that were differentially expressed between experimental groups [121]. To elucidate the potential functional roles of *H. vulgare* siRNAs, Gene Ontology (GO) annotations and protein information for siRNA target genes identified via degradome analysis were retrieved via UniProt identifiers[122]. Enrichment and visualization of GO categories and protein interactions were performed via g:Profiler and Cytoscape version 3.10.1. Graphical representations of the results were generated with the ggplot2 package in R [123–125].

### RT‒qPCR quantification

To validate the small RNA sequencing results, reverse transcription quantitative PCR (RT‒ qPCR) analysis was conducted[126]. Primers targeting mature siRNA sequences were designed via the miRprimer software, and the sequences are provided in Supplementary Table 2 [127]. Stable siRNAs exhibiting consistent expression across samples were selected as reference genes via the RefFinder tool [128]. cDNA synthesis of siRNAs was performed via the Mir-X™ miRNA First-Strand Synthesis Kit (Takara Bio, Kusatsu, Japan) with 400 ng of input RNA, following the manufacturer’s instructions. RT‒qPCR was performed via the FastStart Essential DNA Green Master Kit (Roche Diagnostics GmbH, Mannheim, Germany) on a LightCycler 96 thermal cycler (Roche, Mannheim, Germany) according to the manufacturer’s standard protocols. Each analysis included three biological replicates and three technical replicates per sample. The relative expression levels of the target siRNAs were calculated via the ΔΔCt method, with HVU_siRNA10720 and HVU_siRNA111994 serving as internal reference controls.

## Supporting information

Level of expression identified siRNA

List of RT-qPCR primer

Expression level of siRNA form RT-qPCR and sRNA-Seq analysis

GO annotation siRNA target genes

Target genes from degradome sequencing

## Author Contributions

Conceptualization, M.P.J.; M.B.; methodology, M.P.J. A.M.; formal analysis, M.P.J.; P.B.; A.M.; data curation, M.P.J.; writing—original draft preparation, M.P.J.; writing—review and editing, M.B.; visualization, M.P.J.; A.M.; supervision, M.B.; funding acquisition, M.P.J. All authors have read and agreed to the published version of the manuscript.

## Acknowledgments

This work was supported by the Preludium 18 project (Project Number: 2019/35/N/NZ9/01046), funded by the National Science Centre Poland.

## Institutional Review Board Statement

Not applicable.

## Informed Consent Statement

Not applicable.

## Data Availability Statement

The data presented in this study are openly available in the NCBI SRA at https://www.ncbi.nlm.nih.gov/bioproject/PRJNA1142182, accessed on 31.07. 2024, Accession: PRJNA1142182.

## Acknowledgments

We would like to thank Magdalena Szechyńska-Hebda for supervision of the Preludium 18 project. We are grateful to the PAN Botanical Garden - Centre for the Preservation of Biological Diversity in Powsin for providing seed material.

