## Supplementary figures and images for "Expression Patterns of Small Interfering RNAs in Germinating Barley (*Hordeum vulgare* L.) Seeds with Age-Induced Differences in Viability"

### Expression level of siRNA form RT-qPCR and sRNA-Seq analysis

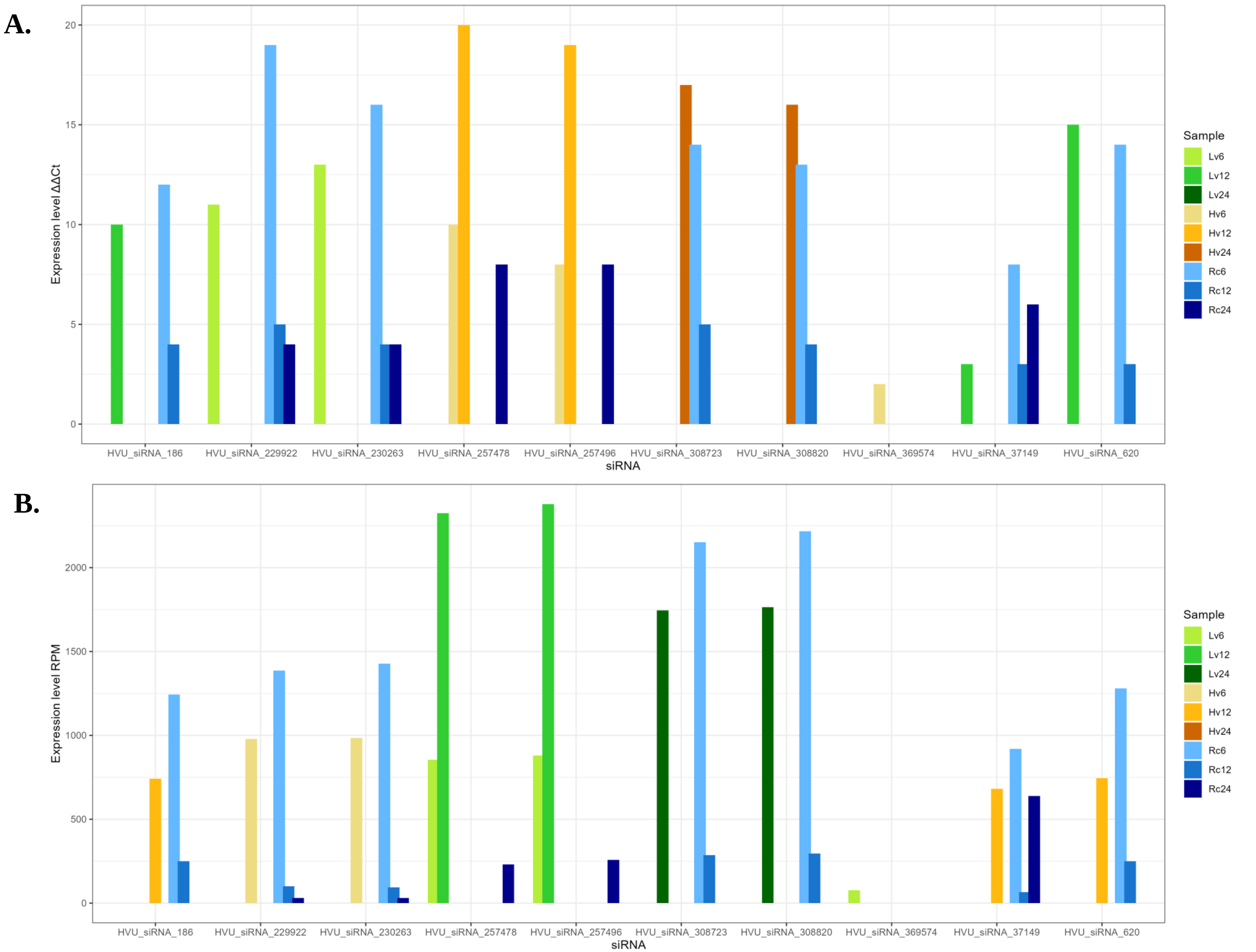

### GO annotation siRNA target genes

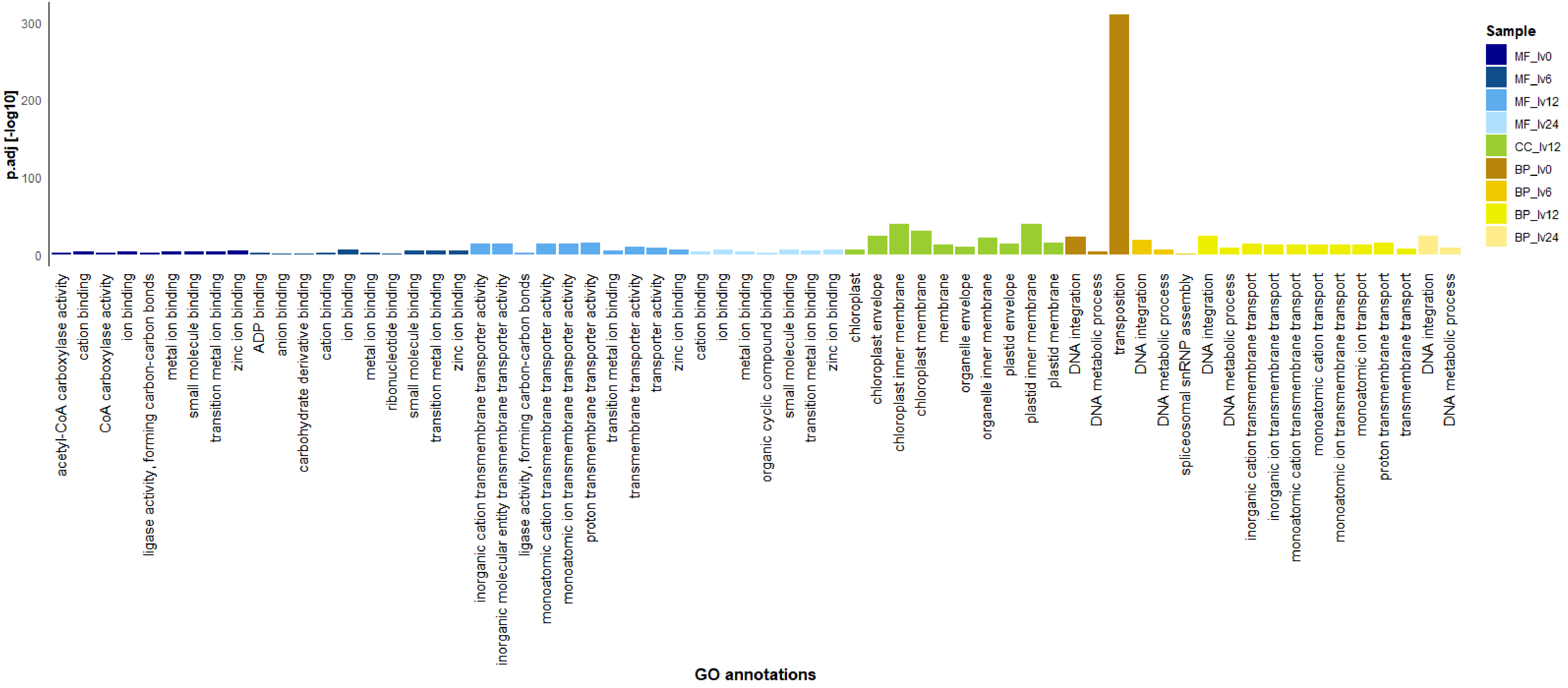

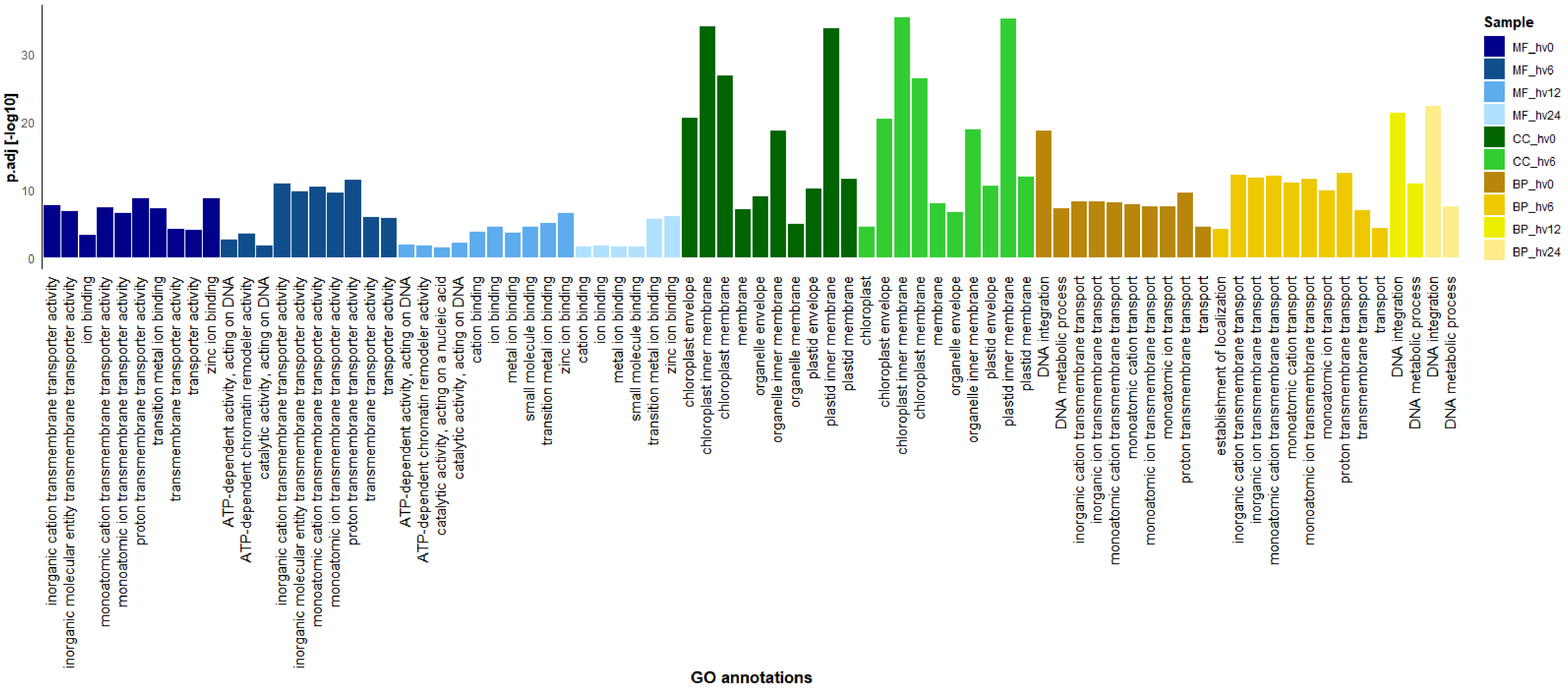

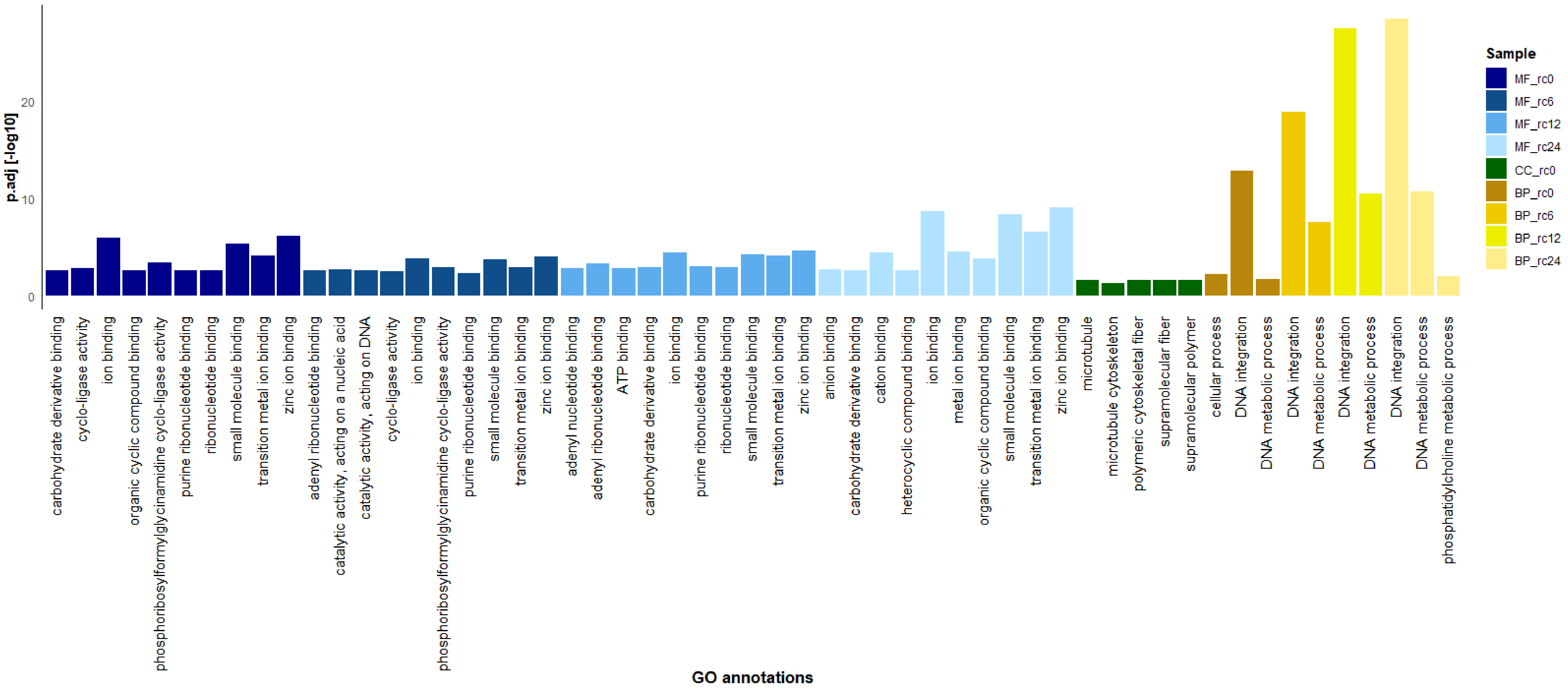
